# Dengue Protease-Mediated Activation and Polarization of Macrophages

**DOI:** 10.1101/2024.11.23.624958

**Authors:** Rajdip Misra, Kaustuv Mukherjee, Joyshree Karmakar, Uttam Pal, Nakul C Maiti

## Abstract

Controlled activation of macrophages to achieve desired phenotypes in vivo and in vitro could lead to possible treatments for several inflammatory and proliferative diseases. This investigation uniquely established that dengue protease enhances the polarization and activation of human and murine macrophages following extracellular exposure. It was observed that macrophages have a robust response to extracellularly administered purified dengue NS2B/NS3 protease; it stimulated the macrophages in classical M1 directions and enhanced both the mitogen-activated protein kinases (MAPK) and c-Jun N-terminal kinases (JNK). The cells exhibited an increase in the intracellular levels of proinflammatory cytokines such as IL-2, IL-6, IL-12, and IFN-γ and an elevation of phosphorylated versions of Akt, P38 MARK, SAPK, and ERK. In tandem, this investigation also established that the heightened responses of stimulated cells were associated with an increase in the cellular reactive oxygen species (ROS) level and the nuclear translocation of the NF-κB (P65) protein. The cell viability assay showed that NS2B/NS3 protease exerts no major toxic effect on macrophages after 24 hours of treatment, even at a dosage (20 μg/ml) which was four times higher than the effective dose (5 μg/ml). Remarkably, we also observed that the native form of the viral protease, which drives its enzyme activities, had no bearing on the antigenic qualities of the enzyme. Thus, our study highlighted the efficacy of dengue viral NS2B/NS3 protease as a non-toxic ex vivo macrophage activating/polarizing agent and may serve a vital role in macrophage-based cell therapy in the near future.

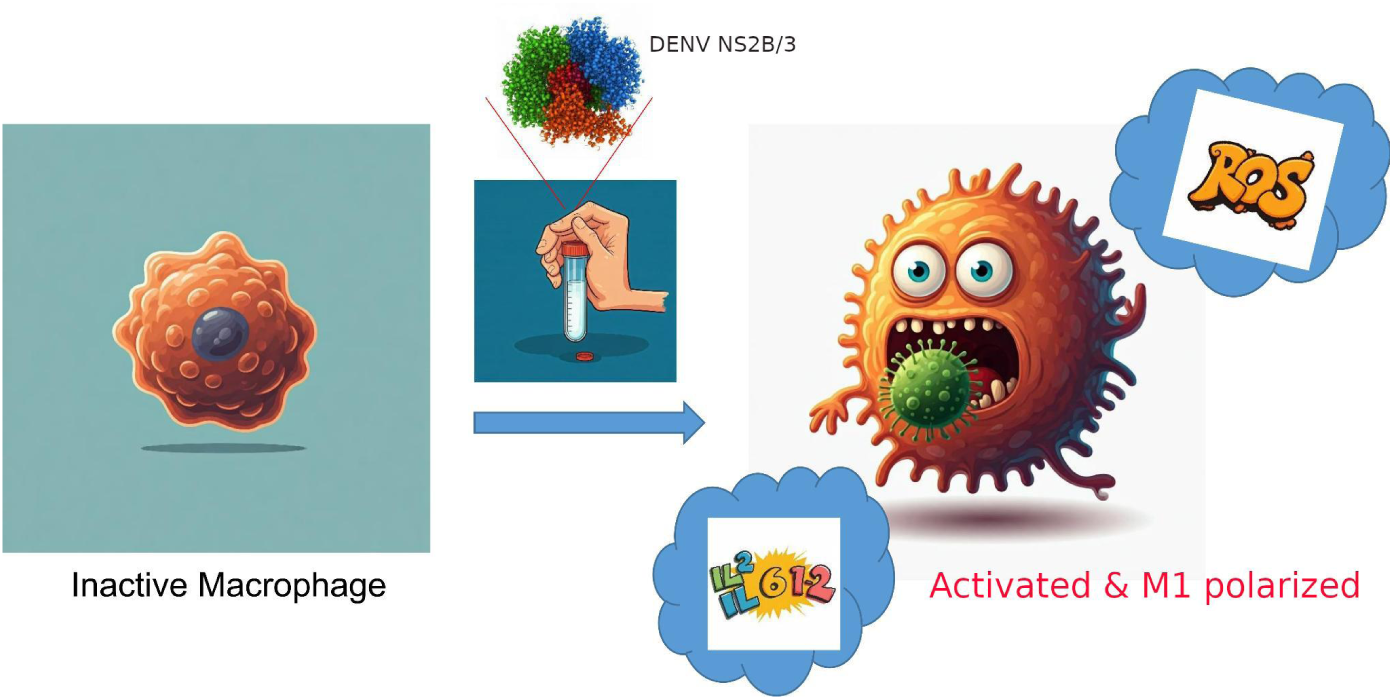

## Introduction

Macrophages are a type of innate immune cell that is specifically located in different tissues throughout the body and provide a crucial homeostatic role in maintaining normal organ function by eliminating endogenous toxic substances such as amyloid beta proteins, dying erythrocytes, apoptotic cells, and surfactants.^1^ Additionally, they are involved with many other biological events, including regulating reactive oxygen species (ROS) endogenous intensities, iron balance, damage to tissue repair, and countless other metabolic functions.^2^ Macrophages, are extensively diverse cells that can rapidly adapt themselves in response to the stimulus arising from local microenvironmental. In vivo, they are very mobile and affect many human illnesses, exhibiting both pathogenic and protective qualities.^3^ Understanding the evolution of macrophage-based cell therapies has focused on its notable capabilities, like promoting tissue regeneration and getting rid of various infections or even cancer cells.^4^

The term “cell therapy” describes the medicinal use of allogeneic or autologous cellular material in a patient.^5^ Cell therapy is still developing today, with continuous research on its efficacy and safety. Cell therapy is an amalgamation of stem cell and non-stem cell treatments that are either unicellular or multicellular. It generally uses autologous or allogeneic cells and may involve genetic editing or formulation changes. It can be administered topically or by injectables or infusions.^6,7^ Cell therapy offers a variety of therapeutic applications, including cancer treatment, immunotherapy, and regenerative medicine. Cell therapies can be categorized by the therapeutic indication they are intended to treat, such as neurological, cardiac, or ophthalmological; by whether they contain cells that are derived from a donor (allogeneic) or taken from the same patient (autologous); or, ideally, by the types of cells, which are frequently categorized by EU regulations.^6^

The efficacy of macrophage-based cell therapy in the traumatic brain injury (TBI) model has been recently highlighted by Kapate and colleagues. A traumatic brain injury is a catastrophic condition caused by rampant inflammation during debris clearance following a regular injury. The disease has no existing treatments other than immediate clinical care. However, when macrophages were programmed with attached discoidal microparticles, encapsulating anti-inflammatory interleukin-4 and dexamethasone, and introduced to the model organisms, lesions were reduced by 56%.^8^ Genetically engineered macrophages have also been used in targeted cancer therapy nowadays. Macrophages’ immunomodulatory potential along with its capacity to invade solid tumors drew attention to their use as immunotherapy. When macrophages were manipulated with genetically engineered anti-human epidermal growth factor receptor-2 (HER2) affibodies onto the extracellular membrane, following further internalization of doxorubicin (DOX)-loaded poly(lactic-co-glycolic acid) nanoparticles (NPs),(NPs), they able to target HER2+ cancer cells and specifically elicit affibody-mediated cell therapy.^9^ Compared to first-generation CAR(Chimeric antigen receptor)-macrophages, the induced pluripotent stem cell-derived macrophages (iMACs) with intracellular toll/IL-1R (TIR) domain-containing CARs showed significantly increased antitumor activity. This approach uplifted the target engulfment capacity of newly engineered iMACs and polarized the macrophages in M1 orientation in a nuclear factor kappa B (NF-κB)-dependent manner. ^10^ Apart from their role in cancer therapeutics, macrophages have also emerged as key regulators of fracture healing, regulating inflammation, reconstruction of tissue, and angiogenesis. Recent breakthroughs in hydrogel-based therapies have created interesting prospects for utilizing macrophages’ modulatory measures to improve fracture repair outcomes.^11^ Among the various macrophage-based cell therapies, ex vivo polarization of macrophages offers several advantages, particularly in the context of research, therapeutic applications, and drug development.^12^

Ex vivo polarization of macrophages is the process of separating macrophages from an organism and stimulating them in a controlled laboratory environment to adopt various functional phenotypes. Macrophages are extremely adaptable cells that can switch between two functional states: M1 or the classically activated triggers the pro-inflammatory responses and M2 or alternatively activated macrophages promote anti-inflammatory responses. In Most of the studies associated with ex vivo polarization and adoptive transfer use M2-polarized macrophages, which minimize inflammation and reportedly improve wound healing. This method has been studied in a wide range of target diseases, including heart tissue and acute kidney damage, spinal cord injury, cutaneous wounds, autoimmune encephalomyelitis, hepatic fibrosis, myocardial infarction and, Achilles tendon rupture.^13–17^ However, M1 polarization has been widely used in cancer cell therapy, repairing liver fibrosis, endometriosis, and activating the adaptive immune system.^10,18–20^

Several chemicals may alter macrophage polarization, in addition to the conventional stimulators IL-4, IL-13, and IL-10. However, many of these factors are also associated with additional side effects, making them less ideal for long-term therapy.^21^ There is a continuous endeavor to find new and more efficient materials for targeted activation.^22–24^ In the current study, we evaluated the effectiveness of a non-structural protein of viral origin as an ex vivo polarizing/activating agent for macrophages. Specifically, we have studied the dengue viral serine protease (NS2B/NS3) exposure to both the human and murine macrophages. The dengue virus (DENV) trypsin-like serine protease, NS2B/NS3, which is largely involved in the cleavage of viral polyprotein into several active proteins, also contains multiple immunogenic motifs that trigger the adaptive immune response in dengue patients.^25^ To establish the polarizing effect of this biomaterial, expression of several phosphorylated signaling molecules like Akt Ser473, ERK, JNK/SAPK, and p-P38 MARK; intracellular and extracellular cytokine levels, levels of type II IFN as well as pro-inflammatory cytokines such as IL-1 and IL-6; cellular reactive oxygen species (ROS), etc. were thoroughly investigated.

## Results

### NS2B/NS3 shows minimal toxicity to murine macrophage J774.A1

Cell toxicity is one of the first endpoints evaluated in drug development and presents valuable data for establishing experimental parameters and implementing doses for in vitro pharmacology studies. Charged molecules have been shown to trigger cell membrane permeabilization, and, at certain doses or concentrations, they impose cytotoxicity. It is necessary to check the toxic effect on cells upon treatment with any foreign particle. To check the toxicity related to purified dengue NS2B/NS3 protease on macrophages, cells (1 × 10^4^ /ml/well) were treated with different dosages (untreated/buffer control,0.5,1,2,5,10 & 20 µg/ml) of the protein, and a cell viability assay was conducted. The result from the viability assay showed that NS2B/NS3 protease didn’t cause major toxicity to macrophages after 24 hours of treatment (Fig. 1). Even at the highest dosage of 20 μg/ml, cell mortality was not observed at 24 hours of treatment. In 48 hours of treatment, the protease showed no significant toxicity to cells up to 10 μg/ml. At further higher doses (20 µg/ml), very slight cell death was observed, possibly due to exposure to a higher stimulant concentration for longer period. As the protein showed no significant toxicity to the cells up to a dosage of 10 µg/ml, even after 48 hours of treatment, we used a maximum dosage not exceeding 10 µg/ml for our assays.

**Figure 1:**
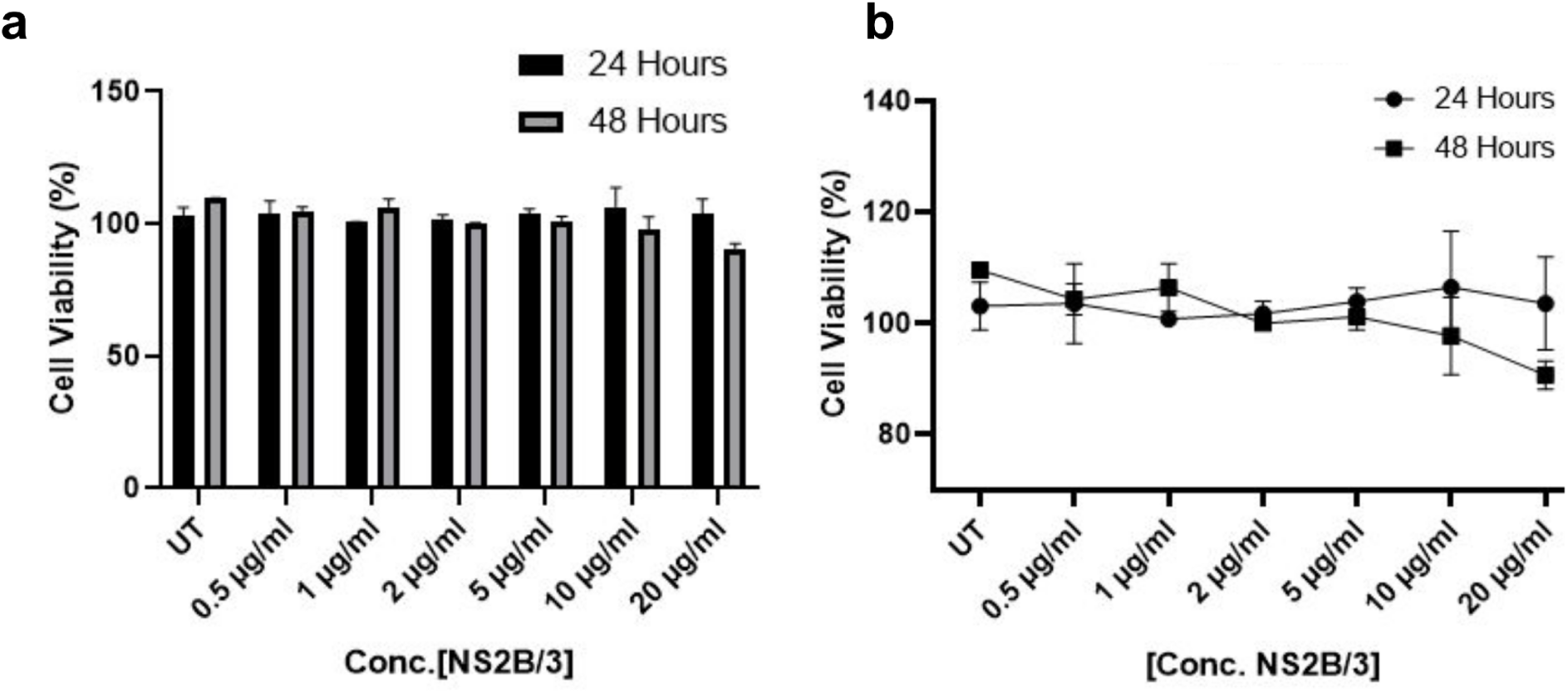
Cell viability assay in Murine Macrophage J774.A1 upon treatment with various doses (μg/ml) of NS2B/NS3 protease for 24 and 48 hours. a. The bar graph represents a non-toxic effect of NS2B/NS3 protease on macrophages at 24 hours of treatment, even at a 20 μg/ml dose. After 48 hours of treatment, no cell death was observed at a dose of up to 5 μg/ml. b. Relative changes in cell viability with different dosages of NS2B/3.

### NS2B/NS3 testament elevates intracellular pro-inflammatory cytokines

So far, we observed that dengue NS2B/NS3 possesses minimal toxicity to murine macrophage J774.A1. Following the cytotoxicity assay, experiments were carried out to determine the efficacy of the dengue serine protease in stimulating cell activation and polarization. Firstly, the intracellular pro and anti-inflammatory cytokine levels of murine macrophage J77A.1 (1 × 10^6^/ml/well), upon exposure to various doses of the protease (1, 5, and 10 µg/ml) for 30 minutes were analyzed by flow cytometry. The results showed a significant increase in the intracellular cytokine levels of several pro-inflammatory cytokines, such as IL-2, IL-6, IL-12, and IFN-γ, in treated macrophages than the control (Fig. 2). Among the treatments, cells treated with 5 µg/ml showed more statistical changes than the other doses, with a consistent increase in different pro-inflammatory cytokine levels (Fig. 2).

**Figure 2.**
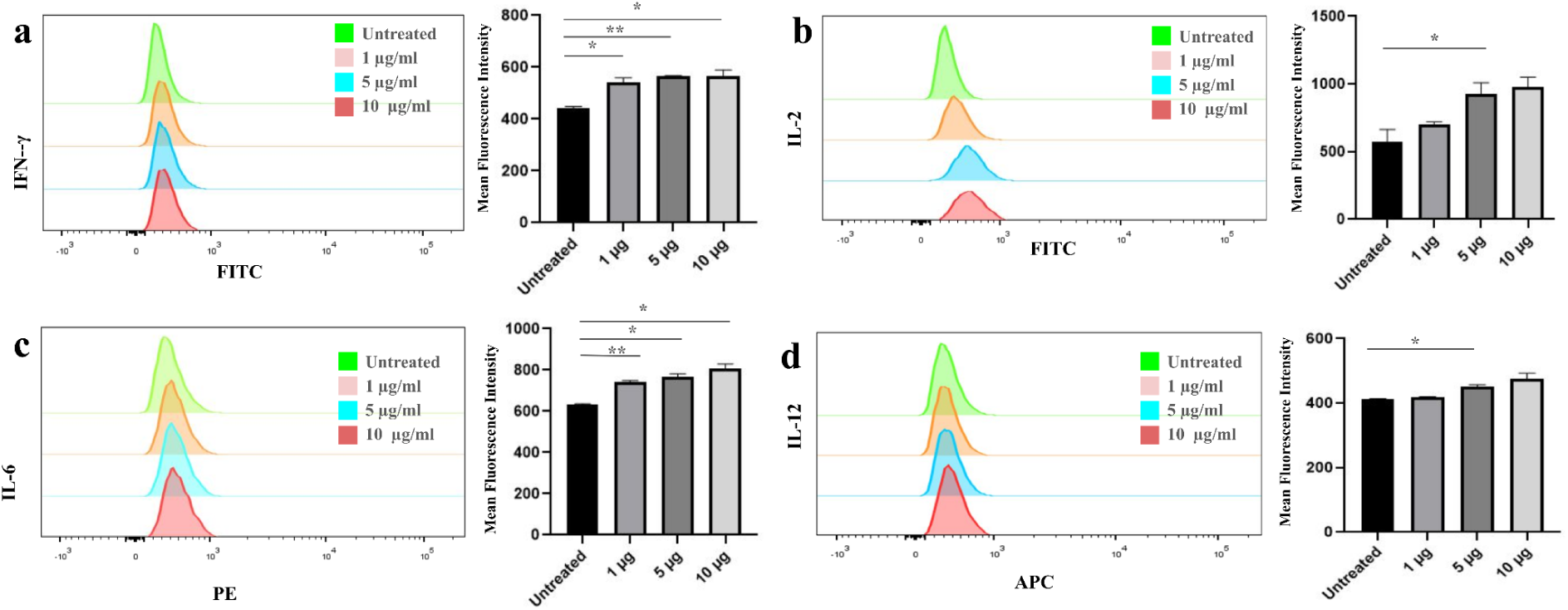
Analysis of intracellular pro-inflammatory cytokine levels in macrophages upon treatment with dengue serine protease by flow cytometry analysis. The results suggest an increase in the expression levels of intracellular pro-inflammatory cytokines upon treatment with different dosages of NS2B/NS3 protease.

### Cellular reactive oxygen species (ROS) levels increases upon exposure to NS2B/3

Furthermore, we quantified the changes in the cellular reactive oxygen species (ROS) in macrophages following treatment with 5 µg/ml NS2B/NS3 protease for 30 minutes. The ROS level is used to confirm pro-inflammatory activation and subsequently M1-polarization of macrophages. ROS plays a significant role in regulating the M1 phenotype, which is majorly involved in the clearance of pathogens.^26^ Macrophages (J774.A1, 1 × 10^6^/ml/well) were scrapped in 1X PBS and incubated with ROS-sensitive dye H2-DCFDA for 30 minutes, and analysis of ROS level was carried out using flow cytometry analysis. The cells treated with dengue NS2B/NS3 protease reportedly exhibited higher ROS generation than the untreated cells (Fig. 3). The histogram suggests a sharp increase in the P4 population from 18.4% in control cells to 36.9% in treated cells.

**Figure 3.**
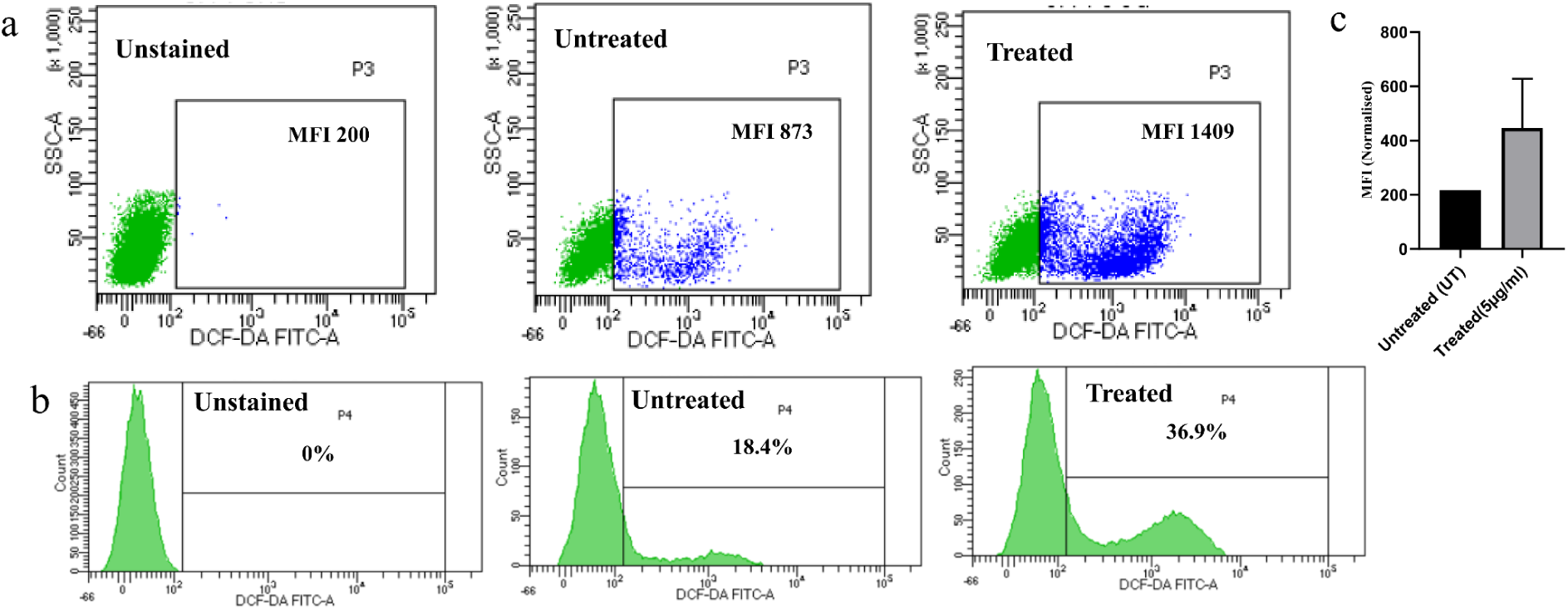
Treatment with NS2B/NS3 effectively raises macrophage reactive oxygen species (ROS) levels. a. The histogram represents a significant increase in the reactive oxygen species levels in the treated cells; b. The bar graph represents the P4 population relative mean fluorescence intensity (MFI) of FITC-H2DCFDA among control and treated cells.

### NS2B/NS3 elevates gene expressions of pro-inflammatory cytokines in Macrophages

Next, we quantified the changes in gene expression levels of both pro and anti-inflammatory cytokines. Macrophages (J774A.1, 1 × 10^6^/ml/well) were exposed to 5 µg/ml NS2B/NS3 protease for 24 hours, later scrapped in 1X PBS and analysis of gene expressions was carried out using the standard protocol.^27^ The gene expressions of a few M1 (IL-12, and TNFα), and M2 markers (Mannose receptor and TGF-β) were analyzed by real-time PCR (Figure 4). We observed a clear change in the gene expression levels of two significant M1 polarization markers, TNFα and IL-12. The fold change in the TNFα gene upon treatment with NS2B/NS3 was increased by more than four in treated cells than in untreated (Fig. 4). Gene expression in IL-12 was also doubled in the treated cells than in the control cells. Though the M1 markers showed a clear upregulation upon treatment, downregulation was observed in the gene expression of anti-inflammatory markers (Mannose receptor, and TGF-β). The expression levels of TGF-β in untreated cells were slightly reduced in treated cells. Apart from TGF-β, gene expression of mannose receptor, another M2 polarization marker, remained more or less similar compared to untreated cells. The collective analysis from the above assays suggested an M1 polarization of macrophages upon treatment with dengue protease.

**Figure 4.**
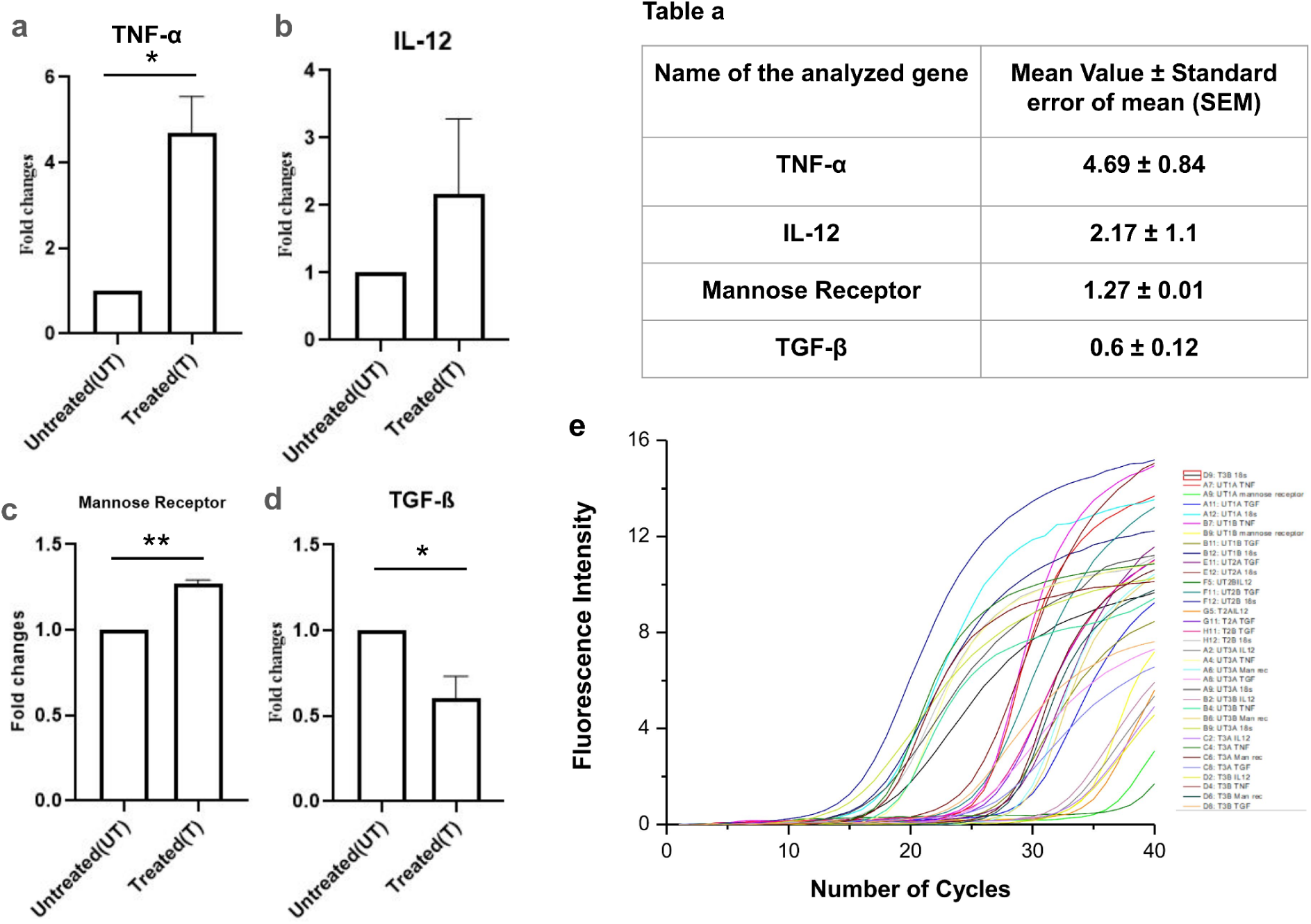
Genetic expression analysis by Real-Time PCR. a-b.The genetic expression of M1 polarization markers such as TNFα and IL-12 were elevated upon treatment with dengue protease; c-d. The genetic expression of M2 polarization markers such as Mannose receptor, and TGF-β were more less remained similar or slightly reduced upon treatment; e. The figure represents the melting curves of PCR cycles; Table A displays the mean value of fold changes along with SEM values.

### NS2B/NS3 evokes elevated MAPK and Akt signaling in macrophages

As of now, we established the effectiveness of dengue NS2B/NS3 protease as an M1 macrophage polarizing agent. Next, we investigated the possible mechanism of action of the protein of our interest. The mitogen-activated protein kinases (MAPK) and Akt, a downstream signaling protein of the phosphatidylinositol 3-kinase (PI3K) pathway, are majorly involved in various physiological conditions in eukaryotes including the regulation of biological activities like apoptosis and cellular growth.^28,29^ Elevated levels of these signaling molecules are also noted during infections. Certain flaviviral infections, such as DENV and Japanese Encephalitis (JE), have been shown to elevate Akt phosphorylation at the Serine 473 (Ser 473) residue.^28^ Along with Akt, the serine/threonine-specific protein kinases regulate various cellular events, including gene expression, proliferation, cellular differentiation, apoptosis, and cell survivability.^29^ A few important members of the MAPK family include ERK 1/2, JNK 1/2, p38 MAPK, etc.^30^ As both the Akt Ser 473 phosphorylation and several MAPKs signaling play key roles in cellular homeostasis, we investigated the possibility that the extracellular NS2B/NS3 protease could affect the cell’s MAPK and Akt expression, leading to the downstream activation and M1 polarization of macrophages. The expression levels of different phosphoproteins in the MAPK family pathway, such as p-P38 MAPK, p-ERK-1/2, and p-JNK/SAPK in murine macrophages (J774.A1, 1 × 10^6^/ml/well) following 24-hour treatment with different doses of NS2B/NS3 protease (UT, 1,5, 10 µg/ml) was checked by western blotting (Fig. 5). Additionally, the total/PAN levels of these protein expressions were also monitored to check for changes in phosphorylation. The ratio of phosphorylation status and the total protein expression levels were used to assess the relative fold changes of different MAPK and Akt expressions in macrophages. Stimulation with purified NS2B/NS3 protein showed enhanced phosphorylation levels of p38 MAPK, JNK/SAPK, ERK1/2 proteins across all protein concentrations. The activatory phosphorylation of Akt at Ser473 was also elevated in stimulated macrophages, particularly upon treatment with 5 µg/ml of protein.

**Figure 5.**
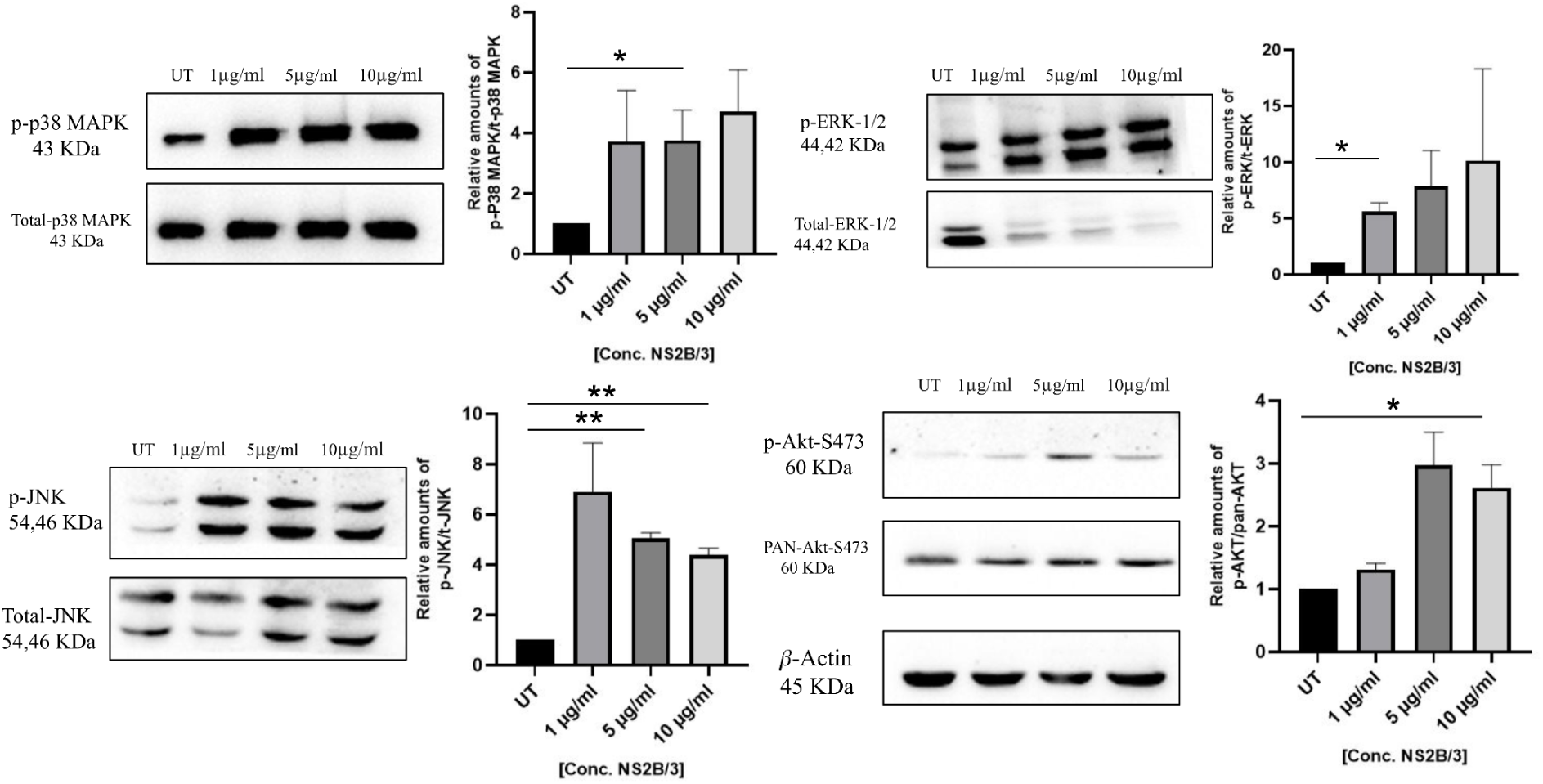
Extracellularly treated dengue NS2B/NS3 protease activates the MAP kinase and Akt expressions in murine macrophages by western blot analysis. The results suggest a sharp increase in the expression levels of several phosphoprotein expressions upon treatment with different dosages of NS2B/NS3 protease.

### NS2B/NS3 protease promotes NF-κB (p65) nuclear translocation

Macrophages responded strongly to NS2B/NS3 protease stimulation, consequently activating cellular responses and M1-polarization with an upstream elevation of MAPK and Akt phosphoproteins. According to the literature, the nuclear translocation of NF-κB protein is associated with several cell activation events, such as ROS, RNS generation, cytokine secretion, and inflammation in macrophages.^31^ Apart from that, NF-κB plays a crucial role in coordinating an organism’s or cell’s reactions to various stress or dangers. It can therefore be triggered by a wide range of stimuli, including physical stressors like heat^32^, cold^33^, ionizing or UV radiation^34^, mechanical shear pressure^35^, and pathogens like bacteria^36^, viruses^37^, and even parasites^38^. To observe the mechanism of action of NS2B/NS3 protease on classically M1-polarized macrophages following an upstream MAPK pathway activation, changes in intracellular localization of NF-κB protein were monitored by both confocal microscopy and western blot analysis. For confocal microscopy, both the J774.A1 (murine macrophage) and THP-1 (human macrophages) cells (1 × 10^5^/ml) were stimulated with 5 µg/ml of purified active NS2B/NS3 protease and incubated for 1 & 2 hours, respectively. Fixed cells were stained with the anti-NF-κB p65 subunit antibody and the respective fluorophore-conjugated secondary antibody before examination by microscopy. The resulting images from the said experiments are pictured in Figure 6. Compared to untreated cells, stimulated macrophages show clear differences in the cellular localization of the NF-κB p65 protein. The signal of the protein (red) evidently increases in the nuclear compartment and is seen to overlap with the DAPI signal (blue) in treated cells. In untreated cells, the NF-κB p65 signal remains distributed throughout the cell (Fig. 6). Our microscopic observations of the NF-κB nuclear translocation were additionally supported by western blot analysis (Fig. S1). Both analyses clarified a significant translocation of NF-κB protein from cytosol to the nucleus in the NS2B/NS3 treated cells.

**Figure 6.**
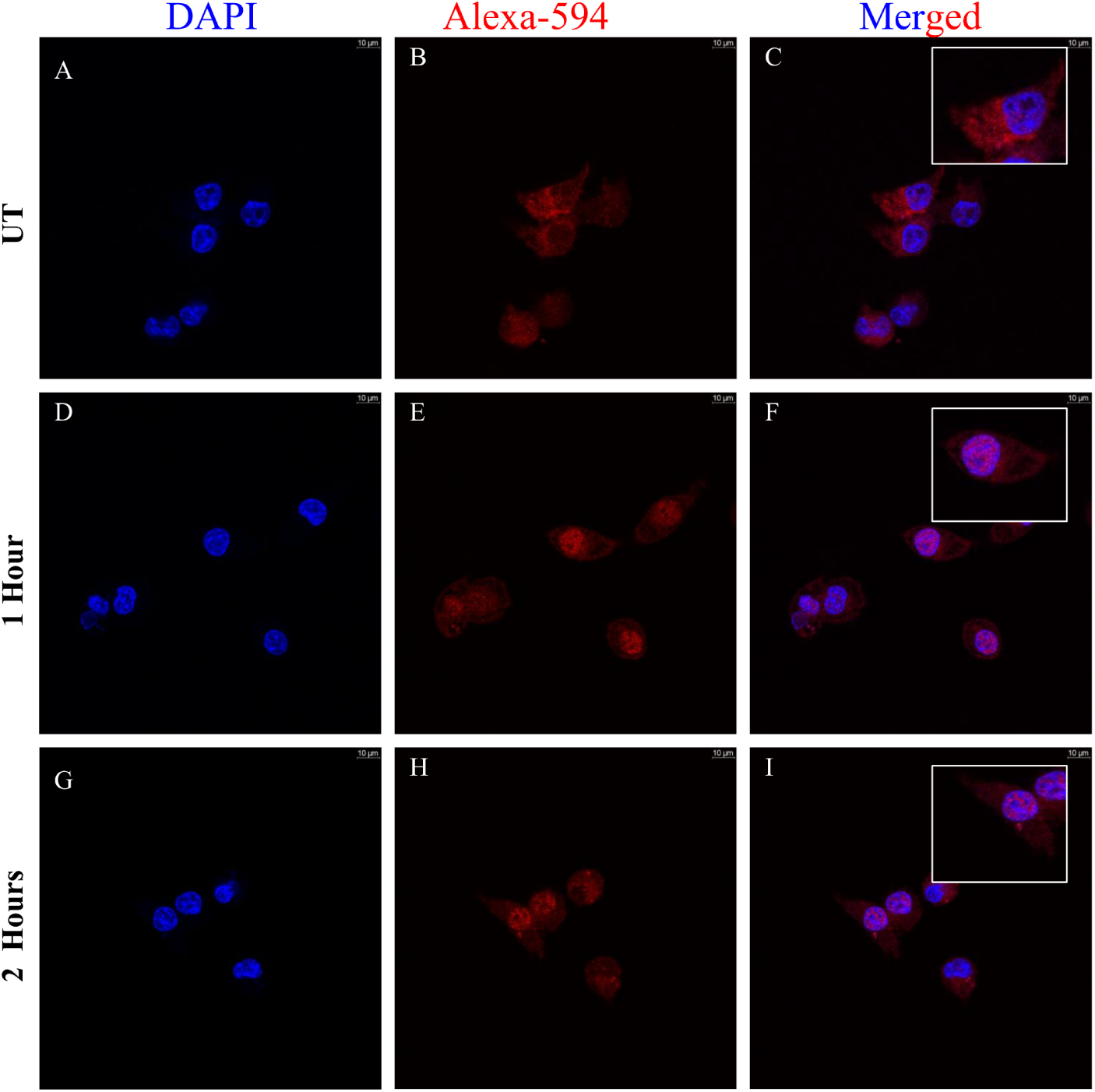
Nuclear translocation of the NF-κB protein of Murine Macrophage J774.A1 upon treatment with 5 μg/ml of NS2B/NS3 protease for 1 and 2 hours, respectively. The nuclear translocation of NF-κB is a significant marker of cell activation and M1 polarization in macrophages. The left panel (A, D, and G) shows cells stained with DAPI. The middle panel shows cells stained with Alexa 594 conjugated anti-NF-κB antibody (B, E, and H). The right panel shows the colocalization of the different dyes (C, F, and I).

### Heat-denatured NS2B/NS3 can also activate macrophages

So far, we have observed that extracellularly given purified dengue NS2B/NS3 protease can activate immunological signaling in murine and human macrophages. We next checked if the inactivated form of the enzyme, devoid of its enzyme activities, could stimulate the innate immunity of murine and human macrophages. The protein, once purified and decontaminated, was boiled at 300 for 10 minutes for denaturation, and the distortion of its native structure as well as enzyme activities were confirmed by circular dichroism spectroscopy (Fig. 7a) and an enzyme assay (Fig. 7b), respectively. Macrophages (J774.A1 & THP-1) were treated with distorted and inactivated 5 µg/ml NS2B/NS3 protease, and nuclear translocation of NF-κB protein was monitored by confocal microscopy as before. The cellular localizations of NF-κB p65 subunit in treated murine macrophages (Fig. 8) and human macrophages (Fig. 9) were compared. The results from the confocal microscopy suggest that heat-inactivated NS2B/NS3 can still stimulate the innate responses in both murine and human macrophages compared to the stimulation with the active NS2B/NS3 protein. Almost no significant change in the translocation of the NF-κB between macrophages treated with heat-treated or native protein was observed. This experiment highlights that the wild-type structure of NS2B/NS3 protein, which powers its enzymatic activities, is not involved in its ability to activate macrophage immune responses.

**Figure 7.**
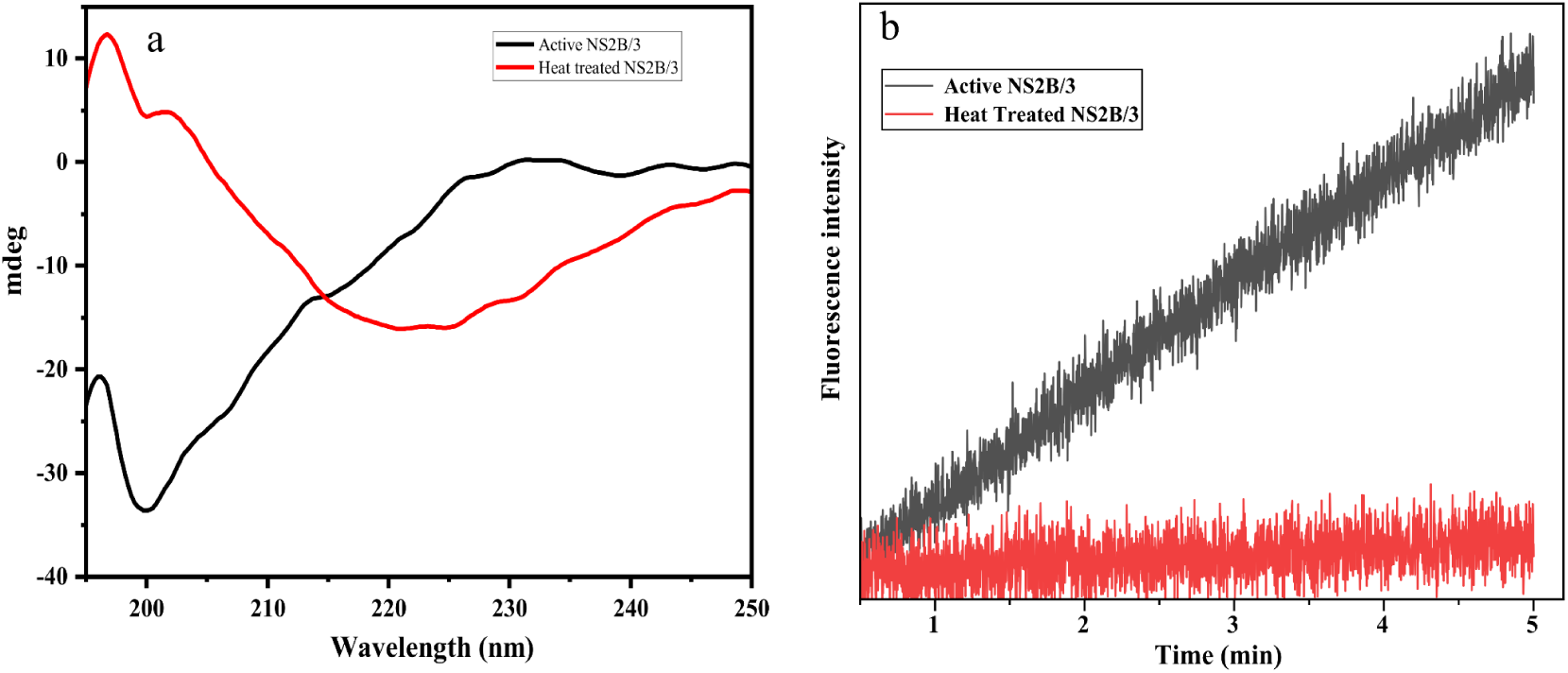
Effect of heat treatment in the structure and function of NS2B/3. a. Effect of heat treatment in the structure of NS2B/3. The heat treatment changes the secondary structure of the protein from more random coil confirmation to a more stable alpha-helical propensity. b. Kinetics Assay: Heat treatment demolishes the enzyme activity of NS2B/3.

**Figure 8.**
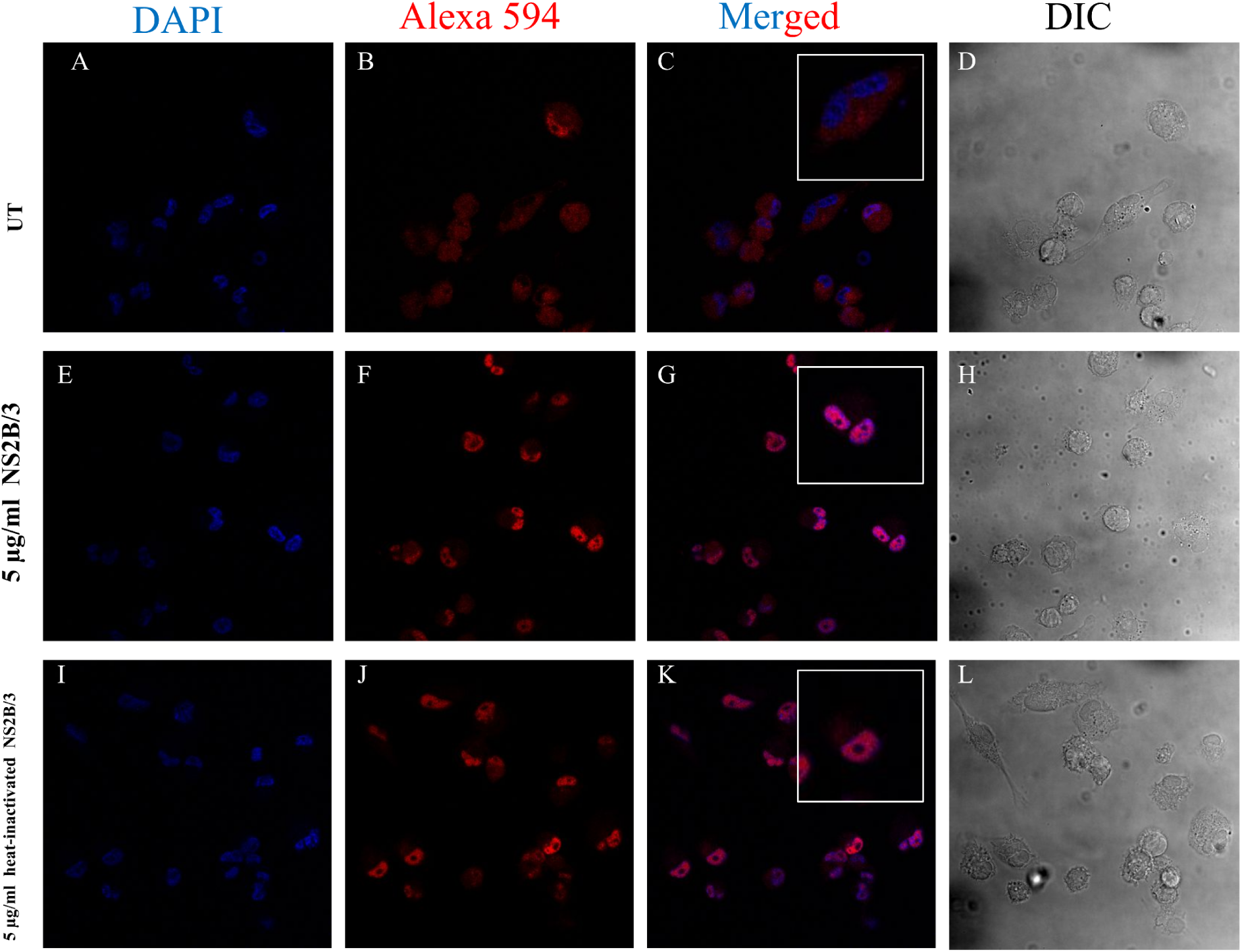
NF-κB nuclear translocation in murine macrophage J774.A1 upon treatment with 5 μg/ml of native and heat-inactivated NS2B/NS3 protease protease for 1 hour. The left panel (A, E, and I) shows cells stained with DAPI. The middle left panel shows cells stained with Alexa 594 conjugated anti-NF-κB antibody (B, F, and J). The middle right panel shows the colocalization of the different dyes (C, G, and K).

**Figure 9.**
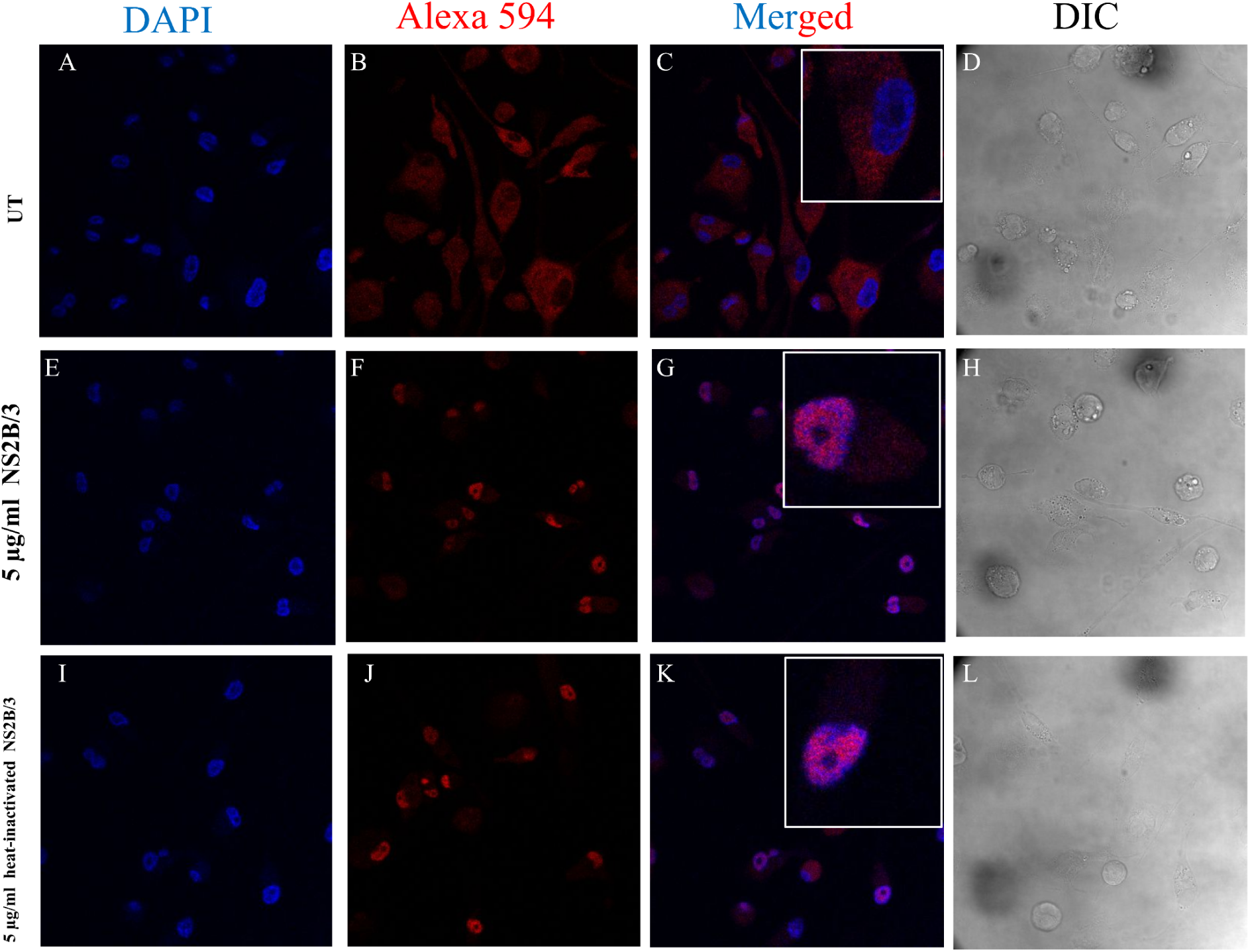
NF-κB nuclear translocation in human macrophage THP-1 upon treatment with 5 μg/ml of native and heat-inactivated NS2B/NS3 protease protease for 1 hour. The left panel (A, E, and I) shows cells stained with DAPI. The middle left panel shows cells stained with Alexa 594 conjugated anti-NF-κB antibody (B, F, and J). The middle right panel shows the colocalization of the different dyes (C, G, and K).

## Discussion

### Ex vivo treatment with dengue serine protease promotes M1-polarization

Ex vivo macrophage polarizing agents are substances or compounds that alter the differentiation and functional polarization of macrophages outside of a living body. Macrophages are important immune system components and can be polarized into different states based on the signals they receive. This polarization can affect their behavioral patterns and their ability to promote or resolve inflammation. The elevation of high levels of proinflammatory cytokines can classify the M1 polarization. The M1-polarized macrophage has been widely used in the treatment of various diseases. According to the research, macrophages polarized in the M1 orientation performed a stronger therapeutic response in the treatment of liver fibrosis.^18^ Not M2, but M1-polarized macrophages effectively increased the recruitment of endogenous macrophages into the liver that has undergone fibrosis. Apart from their protective roles, M1-directed macrophages also enhanced the number of activated natural killer cells (NK cells) in the fibrotic liver. M1 polarization also reportedly increases the phagocytic and tumor-killing abilities of macrophages. The lipopolysaccharide-induced M1 macrophages successfully annihilate HER2-positive tumor cells in vitro and in vivo, showing adverse therapeutic effects against solid tumors.^19^

In our study, cells responded strongly to the extracellularly administered NS2B/NS3 at a dosage of 1μg/ml–10μg/ml, optimizing at 5μg/ml. Classically activated macrophages, additionally referred to as M1 macrophages, serve a substantial part in the immune response to infections and inflammatory events. M1 polarization is distinguished by the production of an array of pro-inflammatory cytokines such as IL-6, IL-12, IFN-γ, and TNF-α.^19^ We observed a similar trend where treatment with the protease strongly uplifted the intracellular IL-2, IL-6, IL-12, and IFN-γ levels in murine macrophages (Fig. 2). Reportedly, 5 μg/ml of NS2B/NS3 also increased the secretory IFN-γ in murine macrophages (Fig. S2). Another parameter to check the classical M1 activation in macrophages is to observe the cellular reactive oxygen species level (ROS).^26^ The cells treated with dengue NS2B/NS3 protease reportedly also exhibited higher ROS generation than the untreated cells (Fig. 3). All these experimental data were further supported by the gene expression analysis of pro and anti-inflammatory cytokines. Both the M1 polarization markers, TNFα and IL-12 were elevated by several folds in the treated cells compared to untreated cells (Fig. 4a & 4b). The treatment with 5 μg/ml of NS2B/NS3 however didn’t increase the gene expressions in M2 polarization markers. Instead, gene expression of TGF-β was reduced by more than 40% in the treated cells than in the untreated. Collectively, it was evident that the protease treatment could stimulate the classical M1 polarization and activation in macrophages.

### NS2B/NS3 is relatively non-toxic to use as a polarizing agent

Among the various molecules used as macrophage polarization agents, two of the most frequently used are lipopolysaccharide (LPS) and graphene oxide (GO). According to global researchers, lipopolysaccharide (LPS), an important outer membrane component of gram-negative bacteria, is associated with severe cytotoxicity. 10 μg/ml LPS decreases cell viability by 50% in human corneal epithelial (HCE) cells within 24 hours of treatment. Further high dosages (>10 μg/ml) resulted in extensive cell death.^39^ LPS also induces cells to secrete cytokines that correspond to lethal toxicity in mice.^40^ Another chemical, graphene oxide (GO), is now employed as a macrophage-polarizing compound and has also been linked to significant cytotoxicity. GO and graphene-based nanoparticles (GPN) can interact with DNA, potentially causing damage.^41^ Aside from that, 100 nm GO reduced cell viability and proliferation, induced extensive lactate dehydrogenase (LDH) leakage, and produced ROS in Leydig (TM3) and Sertoli (TM4) cells.^42^ Again, graphene oxide (GO), few-layer graphene (FLG), and small FLG (sFLG) were also found to be toxic to normal human bronchial epithelial (NHBE) cells. Reports suggest that the compounds could induce apoptosis and necrosis in the NHBE cells following 6–24 hours of exposure, with cell death reaching 90% after a 5 µg/ml dose.^43^ Here we have investigated the potential efficacy of the dengue viral serine protease NS2B/NS3 to induce, activate, and polarize macrophages. The cytotoxicity assay (Figure 1) confirmed that the protein of our interest (NS2B/3) used for macrophages’ activation and polarization purposes, shows negligible toxicity to macrophages. The assay shows that no cell death was observed at 24 hours of treatment, even at the dosage of 20 μg/ml. At 48 hours, no cell death was observed until the 10 μg/ml dosage. At the highest dosage (20 μg/ml) used for the assay, the cell viability observed was >90% (Fig. 1). Based on the toxicity assay, we have used a maximum dosage of 10 μg/ml and a maximum exposure time of 24 hours for future assays, at which no cytotoxicity was observed.

### NS2B/NS3 shows strong antigenic properties elevating MAPK, Akt, and NF-κB signaling pathways

To understand the mechanism of action of dengue NS2B/3, we checked for the phosphorylation of different key signaling proteins from the MAPK and Akt pathways. Results from the western blot analysis (Figure 5) revealed a several fold increase in the phosphorylation level of cellular kinases (P38 MAPK, JNK/SAPK, and ERK) in the treated cells. Following treatment with NS2B/3, Akt serine 473 phosphorylation, associated with several intracellular events like cell survival and apoptosis, also significantly rose.^44,45^ Akt ser473 phosphorylation also plays a significant role in various viral infections. Cases suggest that inhibition of phosphorylation of ser 473 repeatedly decreases the viral titer of DENV in the Huh7 cell line.^46^ Chen et al. additionally discovered the important role of Akt/PI-3K signaling in dengue illness. According to their research, inhibition of the Akt pathway by AR-12, a derivative of celecoxib, effectively prevented the process of replication in hemorrhagic fever-causing viruses.^47^ The increase in Akt ser473 phosphorylation, particularly at a dosage of 5 μg/ml in macrophages upon treatment with NS2B/NS3 (Fig. 5), suggests an increase in the growth signaling of the cells. Additionally, the elevation in the phosphorylation of several other MAP kinases such as p-p38 MAPK, p-ERK, p-JNK/SAPK proteins suggests a strong immunological activation at the cellular level of macrophages. Furthermore, the above observations were also supported by the downstream activation and nuclear translocation of the NF-κB P65 protein. Compared to the control cells, the signal of the p65 protein (red) evidently increases in the nuclear compartment and is seen to overlap with the DAPI signal (blue) in treated cells (Fig. 6) A similar trend was also observed from western blot analysis of p65 protein (Fig. S). Both The activation and translocation of NF-κB protein is a key step in the M1 polarization of macrophages that increase ROS production, promote phagocytosis, and elevate pro-inflammatory cytokines such as IL-2, IL-6, IFN-γ, and TNF-α.^48–51^

### The macrophage activation by Dengue NS2B/NS3 protease is independent of its native structure & enzyme activity

Finally, we investigated the relationship between the native structure that drives the enzymatic activity and the antigenic propensity of the viral protease. NS2B/3, the dengue viral serine protease, is associated with the cleavage of viral polyprotein in the host cell. Using an active enzyme, in particular, protease as an innate boosting agent can be challenging. Proteases are enzymes that break down peptide links between amino acids in proteins and play important roles in various biological processes, including protein digestion, cell communication, and mortality^52^. Using proteases as immunogens can damage cells, specifically if they are not properly controlled or targeted. Due to their proteolytic activities, proteases may also cleave cell surface proteins, resulting in dysregulation of how cells function^53^. Next, we addressed whether the entire structure of the protein is essential for the M1 activation and polarization of macrophages. Heat treatment of NS2B/NS3 changed the secondary structure of NS2B/NS3 from a more random coil propensity to a stable alpha helical conformation (Fig. 6). The change in it’s secondary structure was linked to its enzymatic activities, as one can observe upon heat treatment, the enzyme completely lost its enzyme activity (Fig. 6). However, When the distorted NS2B/3, devoid of any enzymatic activities, was introduced to both murine and human macrophages; it still could stimulate the nuclear translocation of the NF-κB protein and initiate the innate responses in both murine and human macrophages (Fig. 7 & Fig. 8). The experimental results suggest that small domains/motifs, rather than the three-dimensional structure of the protein, play a crucial role in M1-polarization and activation of macrophages.

## Conclusion & Future Prospects

In conclusion, we demonstrated and established the efficacy of the dengue viral NS2B/NS3 serine protease as a non-toxic ex vivo M1 polarization and activation agent. The study evaluated that protease treatment effectively uplifted intracellular and intercellular pro-inflammatory cytokines. The treated macrophages exhibited higher cellular reactive oxygen species (ROS) and a significant increase in NF-κB nuclear translocation than control cells. The protease mainly activates the MAPK and Akt pathways upstream, where it elevates the phosphorylation of several kinases such as ERK, Akt, JNK, and p-38 MAPK. NS2B/NS3 showed no significant toxicity towards the macrophages, making it a potential non-toxic polarization and activation agent. Apart from its ability to induce macrophages, NS2B/NS3 is a naturally occurring protein and thus better biocompatible with cells in comparison to synthetic products or prior arts. Our molecule of interest needs not to be maintained in its native structure. The heat-inactivated protein can still activate and polarize the macrophage; thereafter, it requires less handling and care. The NS2B/NS3 dengue protease, therefore, can be a useful protein for macrophage ex vivo polarization/activation and can be used in macrophage-based cell therapy shortly.

## Materials & Methods

### Reagents

Luria Broth, Ampicillin, and Isopropyl-β-D-thiogalactopyranoside (IPTG) were obtained from Hi-Media. Fluorescein isothiocyanate (FITC), Fluo-3-AM, Fura-2-AM, pluronic F, bovine serum albumin (BSA), 4′,6-diamidino-2-phenylindole (DAPI), trypan blue, paraformaldehyde were obtained from Sigma (St. Louis, MO). The mounting medium was from Amersham Biosciences (Uppsala, Sweden); 2′ 7′ - dichlorodihydro fluorescein diacetate chloromethyl ester (CM-H2DCFDA), Alexa Fluor-594 conjugated anti-Rabbit secondary antibody was from Molecular Probes, Thermo Fisher Scientific (OR, USA. All cytokine ELISA kits were obtained from BD Pharmingen and BD Biosciences (San Jose, CA, USA). Anti-6X histidine mouse primary antibody was obtained from GeneTex (GTX33607). Unless indicated otherwise, all other antibodies were from Cell Signaling Technologies (MA, USA). All cell culture medium, fetal calf serum (FCS), and other reagents were purchased from Invitrogen (Thermo Fisher Scientific, Waltham, MA, USA). West-pico-enhanced chemiluminescent substrate (ECL) and BCA assay kit were purchased from Pierce (Thermo Scientific, Waltham, MA, USA). tBOC-Gly-Gly-Arg-AMR, the fluorogenic enzyme substrate, was purchased from BiotechDesk.

### Bacterial Culture, Protein Expression & Purification

Expression and purification of dengue NS2B/NS3 protease were discussed in our previously published article.^52^ In brief, *Escherechia coli* (stain BL-21 DE3), a non-virulent strain, was grown overnight in Luria broth (LB) medium containing ampicillin (100 μg/ml) at 37°C at a continuous shaking rate of 120 rpm. Overnight-grown primary cultures were used for secondary bacterial culture optimized at 37°C for 3 hours. Isopropyl-β-D-thiogalactopyranoside (IPTG) was added at a concentration of 1mM to the medium and cultured for an additional 18 hours at 18°C to ensure the production of our protein of interest. The cells were then centrifuged at 6500 rpm for 10 minutes to obtain pellets. Pellets were harvested by adding lysis buffer [50 mM Tris, 50 mM NaCl, 5% Glycerol, 2mM Imidazole, 5mM β-mercaptoethanol (β-ME), 10mM PMSF, and protein inhibitor cocktail (PIC)], followed by vortexing, sonication, and centrifugation. The ÄKTA-Start Fast Pressure Liquid Chromatography (FPLC) system was used for affinity and size exclusion purification of dengue NS2B/NS3 protease (Fig. S3A). The column was equilibrated with 50 mM Tris pH 8.5, 50 mM NaCl, 5% glycerol, and 20 mM Imidazole buffer and washed with 50 mM Tris pH 8.5, 50 mM NaCl, 5% glycerol, and 40 mM Imidazole buffer. The protein of our interest was eluted with a buffer containing 50 mM Tris pH 8.5, 50 mM NaCl, 5% glycerol, and 100 mM Imidazole solution. The eluted fraction was subjected to overnight dialysis at 4°C to remove any imidazole present in the solution. Following that, the eluted protein solution was then concentrated using 50 ml 10 KDa concentrator tubes (Sigma) to a final volume of 500 ul. Next, the affinity-purified protein fraction was loaded into Superdex HiLoad 16/600 for further purification by size exclusion chromatography (Fig. S3B). The column was extensively washed with double-filtered (0.45 um filter paper, Millipore) distilled water and equilibrated with a buffer containing 50 mM Tris, pH 8.5, and 50 mM NaCl. The protein fraction was eluted with 50 mM Tris, pH 8.5, and 50 mM NaCl buffer and subjected to overnight dialysis at 4°C. Following that, the eluted protein solution was then again concentrated using 50 ml 10 KDa concentrator tubes (Sigma) to a final volume of 500 ul, and its purity was checked by mass spectrometry, SDS-PAGE, and western blot analysis (Fig. S4).

### Endotoxin Removal

Endotoxin contaminants associated with the purified protein fraction were further removed by treating the protein solution with Polymyxin B Agarose beads (P1411, Sigma), as discussed in previously published articles^54–59^. In brief, the solution containing Polymyxin B was initially washed with 1X Phosphate Buffered Saline (PBS) and then added to the protein solution roughly at a ratio of about 1:5, incubated in ice for 2 hours with gentle tapping at times. The mixture was further centrifuged at 2000 rpm for 10 minutes at 4°C. Following that, the supernatant was collected without disturbing the pellets and stored at −80°C until use.

### Mammalian Cell Culture

Murine macrophage (J774.A1) and human monocytic (THP-1) cell lines were obtained from the Cell Repository of National Centre for Cell Science (NCCS), Pune, India. The cell lines were thoroughly cultured at 37°C with 5% CO_2_ in Iscove’s Modified Dulbecco’s Medium (IMDM, Gibco, Invitrogen) and Roswell Park Memorial Institute (RPMI, Sigma) mediums, respectively, supplemented with 10% fetal calf serum (FCS). THP-1-derived macrophages were generated by treating the cells with Phorbol 12-myristate 13-acetate (PMA, 100 ng/mL) for 48 hours and differentiation was monitored by cellular morphology.

### Cell Viability Assay

The MTT assay was performed to determine the cell viability of macrophages and was based on the reduction of 3-(4,5-dimethylthiazolyl-2)-2,5-diphenyltetrazolium bromide (MTT) by mitochondrial dehydrogenase in viable cells to produce a purple formazan product. J774A.1 cells were seeded in a 96-well culture plate at a density of 5× 10^3^ cells/well and cultured in the absence or presence of different doses (Buffer control/UT, 0.5, 1,2,5,10,20 μg/ml) of NS2B/NS3 protease for 24 and 48 hours at 37°C with 5% CO_2_ in a final volume of 200 μl. Following treatment, spent media was discarded, and 100 μl of fresh media containing MTT (1 μg/μl) was added to each well and cultured at 37°C with 5% CO_2_ until the formazan crystals were visualized. The formazan crystals were dissolved in 100 μl of DMSO. Absorbance was read at 550 nm on a multi-well plate reader. The percentage inhibition of cell viability was calculated as a fraction of control.

### Determination of Intracellular Cytokines

J774A.1 cells were seeded at 1 × 10^6^/ml/well and treated with 1, 5, and 10 µg/ml of NS2B/NS3 protease, incubated for 24 hours at 37°C at 5% constant CO_2_. Cells were then washed, permeabilized, and incubated with cytokine-specific rabbit primary and conjugated anti-rabbit secondary antibodies and analyzed by flow cytometry.

### Determination of Secretory IFN-γ Level

J774A.1 cells were activated using PMA (100 ng/mL). Cells (1 × 10^6^) were treated with 1, 5, and 10 µg/ml of NS2B/NS3 protease for 1 hour at 37°C. Infected macrophages were further incubated for 3 hours at 37°C at 5% CO_2_. Culture supernatants were used to estimate secretory cytokine levels of IFN-γ (551866, BD Biosciences) using ELISA kits.

### Detection of Reactive Oxygen Species (ROS)

Macrophages (1 × 10^6^) were treated with 5 µg/ml of NS2B/NS3 protease and incubated for 30 min at 37°C. ROS-sensitive dye-CM-H2DCFDA (20µM, C6827, Molecular Probes, Invitrogen) was added and incubated for an additional 30 min at 37°C. Cells were washed and analyzed by flow cytometry to estimate ROS generation.

### Genetic Expression analysis by Real-Time PCR

Macrophages (1 × 10^6^/ml) were treated with 5 µg/ml of NS2B/NS3 protease and incubated for 24 hours at 37°C at 5% CO_2_. Total RNA from these cells was extracted using the RNeasy mini kit as per the manufacturer’s instructions. First-strand cDNA was synthesized using template mRNA (1 µg) by Reverse transcription following the manufacturer’s protocol. Real-time PCR was performed with specific primers for M1 (IL-6, IL-12, IFNγ, and TNFα), M2 markers as reported earlier^27^ (Arginase, Mannose receptor, IL-10, and TGF-β), obtained from Eurofins Genomics India Pvt. Ltd. using a DyNAmo Flash SYBR Green qPCR Kit. Relative amounts of target mRNA were quantitated using the Light Cycler 480 II instrument, with 18S rRNA as an internal control. Data were analyzed using the OriginPro 8 software and expressed as a fold change compared with untreated control using the comparative cycle threshold (CT) method.^60^

### Immunoblot Analysis

J774A.1 and THP-1 macrophages (1 × 10^6^) were treated with 1, 5, 10 µg/ml of NS2B/NS3 protease and incubated for 24 hours at 37°C at 5% CO_2_. Cells were scraped into Phosphate Buffered Saline (PBS), sonicated in the presence of protease and phosphatase inhibitor cocktail (Calbiochem, Merck-Millipore) and protein was estimated by BCA assay (Pierce, Thermo Scientific). Equal amounts of protein were resolved in SDS-PAGE (10%) gel and transferred onto a PVDF membrane following the wet transfer method (ref). PVDF Membranes were blocked with Bovine serum albumin ((BSA) and incubated overnight with primary antibodies (1:1,000) against different signaling proteins. Then, membranes were washed using Tris-buffered Saline containing 0.1% Tween-20 (TBS-T), incubated with appropriate HRP-conjugated secondary antibodies (1:2000), and visualized using chemiluminescence substrate (Pierce, Thermo Scientific) in a Chemidoc imaging system (BioRad Laboratories, CA, USA). The primary antibodies used include— p-p38 MAPK (#4511), p38 MAPK (#9212), p-Akt Ser473 (#4060), pan-Akt (#4691), p-ERK1/2 (#4377), ERK1/2 (#4695), p-JNK/SAPK (#4668), JNK/SAPK (#9252), β-Actin (#4970), NF-κB Pathway Sampler Kit (#9936) from Cell Signaling Technology and 6X His tag mAb (#RM146) from GeneTex.

Furthermore, cells were suspended in cytosol isolation buffer (10 mM Tris-Cl, 10 mM NaCl, 1.5 mM MgCl2, 1 mM PMSF, 0.05% NP-40, pH 6.8), vortexed to allow partial lysis of membranes, centrifuged at 1,000 g for 5 min to precipitate nuclear fractions and cytosol was collected as supernatant. The pellet was washed with chilled PBS, and incubated in nuclear isolation buffer (20 mM Tris-Cl, 137 mM NaCl, 1 mM CalCl2, 1 mM MgCl2, 1 mM PMSF, 1% NP-40, pH 8.0) for 30 min. Nuclear fraction was collected after centrifuging the suspension at 1,000 g for 5 min at 4°C. Nuclear fractions were similarly resolved via SDS-PAGE and probed by western blotting. Densitometric analysis of bands was performed using ImageJ software and values normalized to Lamin B band intensity have been represented as bar diagrams.

### Analysis of NF-κB Nuclear Translocation by Confocal Microscopy

J774.A1 and THP-1 cells (1 × 10^5^/ml) were seeded in glass coverslips and incubated overnight in IMDM and RPMI mediums, respectively, at 37°C with 5% CO_2_. THP-1 cells were additionally incubated for 48 hours with PMA before treatment. 5 µg/ml of purified active NS2B/NS3 protease was added to the coverslips and incubated for 1 & 2 hours, respectively, to analyze NF-κB translocation experiments. Unbound fractions were removed by washing with PBS. These treated macrophages were finally fixed using 2% paraformaldehyde, permeabilized (PBS containing 0.3% Triton-X-100), blocked (3% FCS in PBS with 0.3% Triton-X-100), and incubated with anti-NF-κB rabbit primary antibodies (1:800; CST 8242) overnight at 4°C. The coverslips were washed with PBS (1X) containing 0.3% Triton-X-100 and counterstained with Alexa Fluor-594 tagged secondary antibodies (1:2000, Invitrogen) in a blocking buffer for 2 hours at room temperature in the dark. Coverslips were washed again, mounted onto slides using a mounting medium containing DAPI, and sealed using nail polish. Images were randomly acquired from multiple fields in Fluoview FV10i (Olympus Life Science) analyzed using Leica Microsystem Software. Again, J774.A1 and THP-1 cells (1 × 10^5) were seeded in glass coverslips and incubated overnight in IMDM and RPMI medium, respectively, at 37°C with 5% CO_2_. THP-1 cells were additionally incubated for 48 hours with PMA before treatment. Cells were treated with 5 µg/ml of wild-type and heat-denatured DENV NS2B/NS3 protein for 1 hour at 37°C with 5% CO_2_. The analysis of NF-κB nuclear translocation was observed using the similar protocol described above.

### Circular Dichroism Spectroscopy

The far UV circular dichroism of the protein was measured at room temperature with a JASCO J-815 spectrometer under continuous N_2_ flow. The freshly purified and endotoxin-free NS2B/NS3 protease was kept in a 10 mM Tris buffer, and the final concentration used was 10 µM. The far UV (190 to 250 nm) spectra of both the active and heat-treated NS2B/NS3 were taken with a path length of 1 mm. For spectra acquisition, the scan speed and bandwidth were set to 50 nm/min and 1 nm, respectively. Each sample was scanned three times, subtracting the buffer background.

### Enzyme Assay

The enzyme activity of the purified active and heat-treated dengue NS2B/NS3 protease was checked using an enzyme assay modified by Kiat. et al^61^ and discussed in our previously published article^52^. In brief, a 100 μl reaction mixture was prepared containing 100 mM Tris buffer pH 9.0, 300 nM purified endotoxin-free NS2B/NS3 protease, and 50 uM fluorogenic peptide enzyme substrate (tBOC-Gly-Gly-Arg-AMR). The reaction mixture was excited at 350 nm, and following that, a continuous fluorescence reading for 5 minutes was measured at 439 nm using an Agilent Cary Eclipse fluorescence spectrometer. The idea was to observe the fluorescence change due to the active enzyme’s cleavage of free AMC molecules from tBOC-Gly-Gly-Arg-AMR.

### Statistical Analysis

The data represented are mean values derived from at least three independent experiments. Statistical analysis was performed using the parametric unpaired/independent Student’s t-test for two groups and more than two groups of samples, with p < 0.05 deemed statistically significant. Error-bars represent mean ± standard error of the mean (SEM) from three independent experiments. Significant differences were set at nsp > 0.05, *p ≤ 0.05, **p ≤ 0.01, ***p ≤ 0.001, and ****p ≤ 0.0001 and analyzed by GraphPad Prism version 8.0.1.

## Data Availability

All the data associated with different experiments are provided either within the manuscript or the supporting information file. The complete workflow is reported in the “Materials and Method” section of the manuscript.

## Associated Contents

### Supporting Information

Details of the plasmid construct; protein purification data (Affinity and size exclusion chromatography); protein purity data (Mass spectra, SDS-PAGE, and western blot); Secretory Cytokine Elisa; NF-κB western blot: Figures S1–SX; Western blot datasheet: Datasheet-1-2.

## Notes

The authors declare no competing financial interest.

## Acknowledgments

R.M., S.K. acknowledges the financial support by the Council of Scientific and Industrial Research (CSIR-NET-SRF). K.M, U.P. and N.C.M. acknowledge the financial support by the Council of Scientific and Industrial Research.

