## Supporting Information for "Dengue Protease-Mediated Activation and Polarization of Macrophages"

### Current Affiliation: Technical Research Centre, S.N. Bose National Centre for Basic Sciences, JD Block, Salt Lake, Kolkata 700095, West Bengal, India

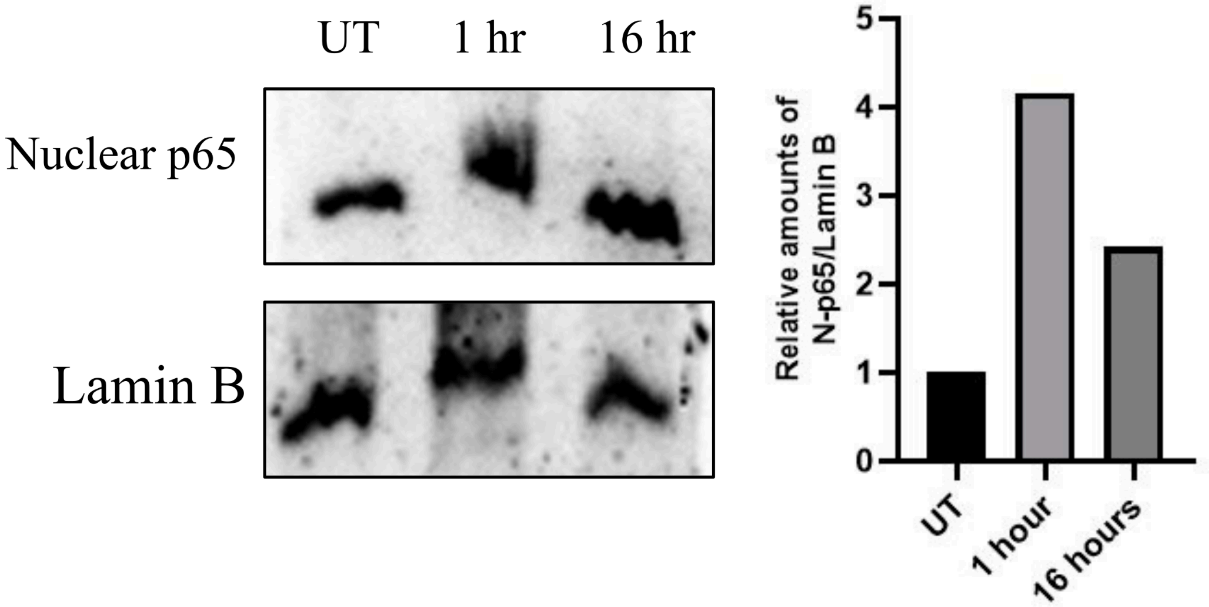


Figure S1. Nuclear translocation of the NF-κB protein of Murine Macrophage J774.A1 upon treatment with 5 μg/ml of NS2B/3 protease for 1 hour by western blot analysis.

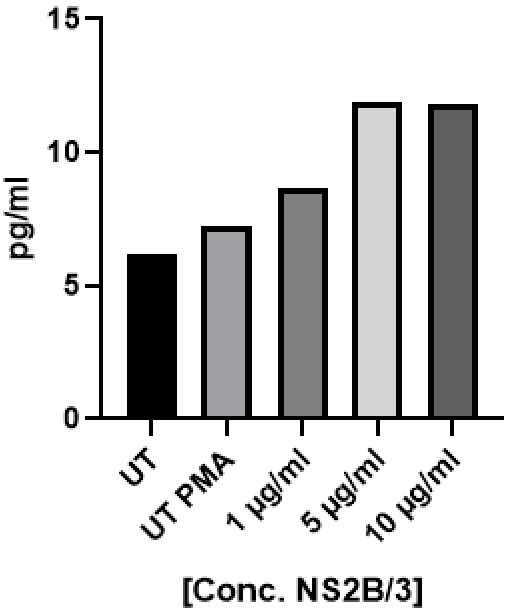


Figure S2. Analysis of the secretory IFN-γ level in murine macrophages upon exposure to 5 µg/ml of NS2B/3.

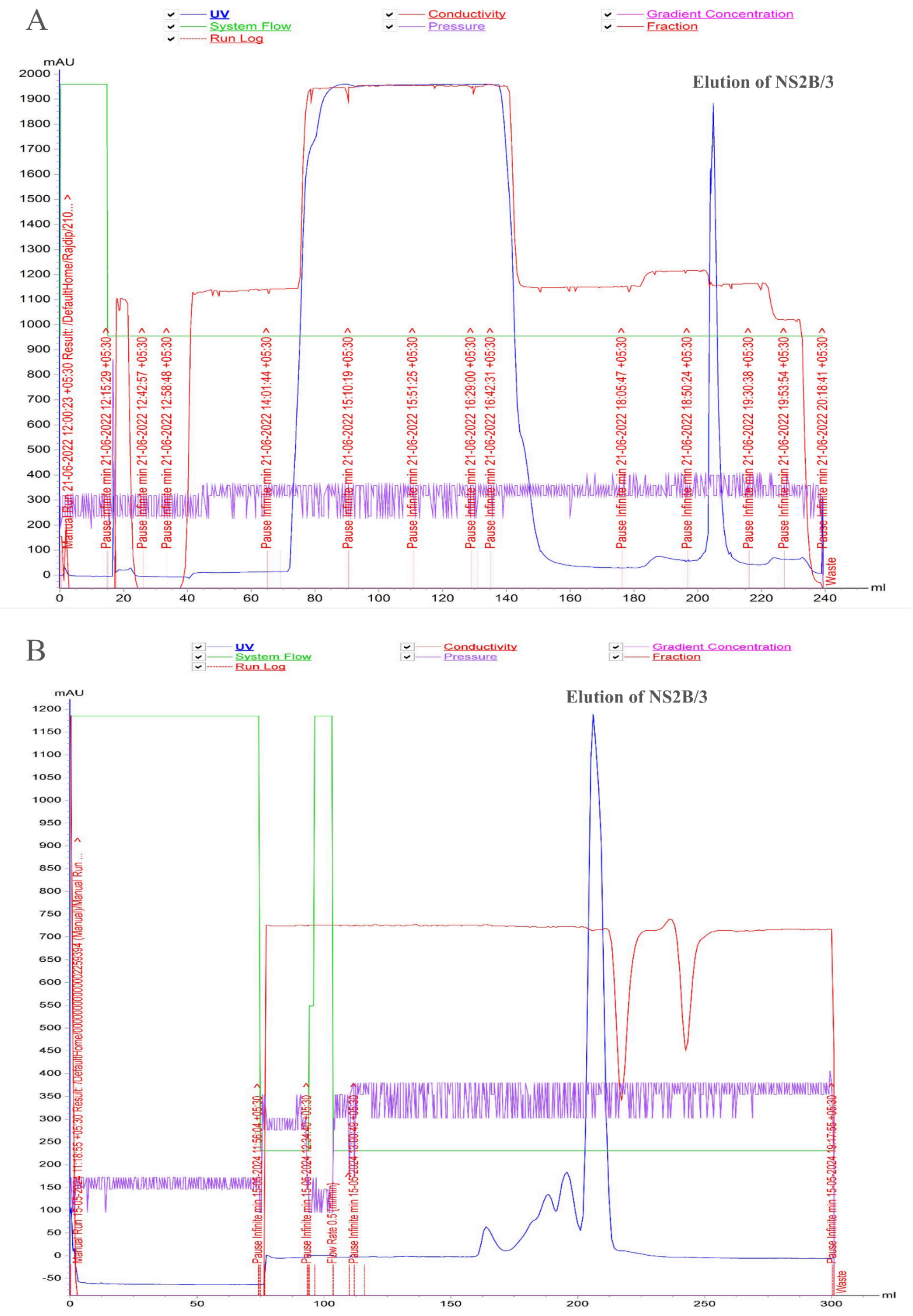


Figure S3. Chromatogram of DENV NS2B/3 purification; A: affinity purification, B. size exclusion purification.

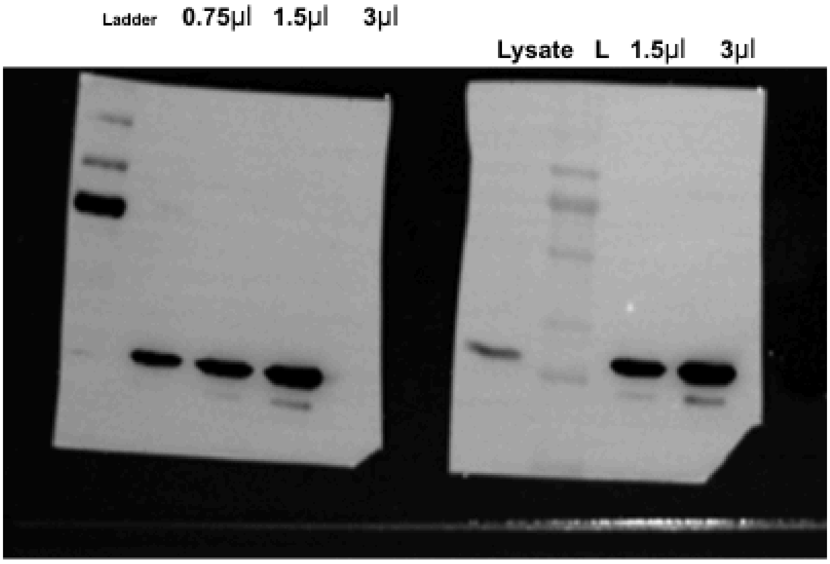


Figure S4. Western blot analysis of dengue NS2B/3.
